# Loss of ASH1L in developing brains causes autistic-like behaviors in a mouse model

**DOI:** 10.1101/2020.11.16.377259

**Authors:** Yuen Gao, Natalia Duque-Wilckens, Mohammad B Aljazi, Yan Wu, Adam J Moeser, George I Mias, Alfred J Robison, Jin He

## Abstract

Autism spectrum disorder (ASD) is a neurodevelopmental disease associated with various gene mutations. Recent genetic and clinical studies report that mutations of the epigenetic gene *ASH1L* are highly associated with human ASD and intellectual disability (ID). However, the causal link between *ASH1L* mutations and ASD/ID remains undetermined. Here we show loss of ASH1L in developing mouse brains is sufficient to cause multiple developmental defects, core autistic-like behaviors, and impaired cognitive memory. Gene expression analyses uncover critical roles of ASH1L in regulating gene expression during neural cell development. Thus, our study establishes a new ASD/ID mouse model revealing the critical function of ASH1L in normal brain development, a causality between *Ash1L* mutations and ASD/ID-like behaviors in mice, and potential molecular mechanisms linking *Ash1L* mutations to brain functional abnormalities.

## Main text

Autism spectrum disorder (ASD) is one of most prevalent neurodevelopmental disorders that have a strong genetic basis (*1*). As hundreds of new ASD risk genes have been identified by recent genetic studies (*2–6*), developing gene-specific knockout animal models has emerged as a priority for further determining whether the identified ASD risk genes are the causative drivers leading to ASD development, as well as for understanding the biological mechanisms underlying the pathogenesis caused by the mutations of ASD risk genes.

ASH1L is one member of Trithorax-group (TxG) proteins that facilitate gene expression through modifying chromatin (*7, 8*). Recent genetic studies on large cohorts of ASD patients reported that mutations of *ASH1L* are highly associated with human ASD (*2–4, 6*). The genetic findings are supported by multiple clinical reports that some children diagnosed with ASD and/or intellectual disability (ID) acquire various disruptive or missense mutations of *ASH1L* (*9–14*). In addition to ASD and ID, patients also display a variety of developmental and behavioral abnormalities including delayed myelination, microcephaly, craniofacial deformity, skeletal abnormality, and feeding difficulties, suggesting critical roles of *ASH1L* in normal embryonic and postnatal development (*9, 11, 12*). However, the causal link between *ASH1L* mutations and clinical ASD/ID remains undetermined. Therefore, in this study we set out to use mice to address: (i) is loss of *Ash1L* in developing brains sufficient to induce ASD/ID-like phenotypes; and (ii) what are the possible molecular mechanisms linking *Ash1L* mutations to the pathogenesis of ASD/ID?

To examine the function of *Ash1L* in mouse development, we generated an *Ash1L* conditional knockout (cKO) mouse line by inserting two LoxP elements into the exon 4-flanking sites at the *Ash1L* gene locus (*Ash1L*^+/2f^). A Cre recombinase-mediated deletion of exon 4 resulted in altered splicing of mRNA creating a premature stop codon before the sequences encoding the first functional AWS (*<u>A</u>*ssociated *<u>W</u>*ith *<u>S</u>*ET) domain. The truncated ASH1L protein contained the N-terminal 1,694 amino acids but lost all functional domains, thus mimicking the disruptive mutations found in patients (Figure 1A, S1A-B). To preclude mouse strain-specific effects on changes of animal phenotypes and behaviors, we backcrossed the wild-type *Ash1l^+/2f^* founders with C57BL/6 mice for more than five generations to reach a pure genetic background. The heterozygous *Ash1L-*KO mice (*Ash1L*^+/1f^) were obtained by crossing the wild-type *Ash1L*^+/2f^ mice with CMV-Cre mice, through which one allele of *Ash1L* gene was deleted in both germlines and somatic cells in progenies. The heterozygous *Ash1L*^+/1f^ x heterozygous *Ash1L*^+/1f^ mating produced normal numbers of embryos. The gross embryos and placentas did not show obvious differences between wild-type and global *Ash1L-*KO (*Ash1L*^1f/1f^) embryos at embryonic day 13.5 (E13.5), and all embryos developed to term with expected mendelian ratios (Figure S1C, S1E), suggesting *Ash1L* was dispensable for mouse embryonic development. The global *Ash1L-*KO newborns displayed similar body size and weight to their wild-type littermates at postnatal day 0 (P0) (Figure S1D). However, without maternal uterine support, all *Ash1L* KO newborns died within 24 hours after birth (Figure 1B, S1E), suggesting *Ash1L* might be critical for establishing and maintaining a stable physiological condition for neonatal survival. Further anatomical analyses did not reveal obvious gross morphological abnormalities of individual organs, except that the majority of *Ash1L-*KO newborns displayed aberrant rib numbers (Figure 1C), which was consistent with the function of TxG proteins in body segmentation and skeletal formation during embryonic development (*15*).

**Figure 1.**
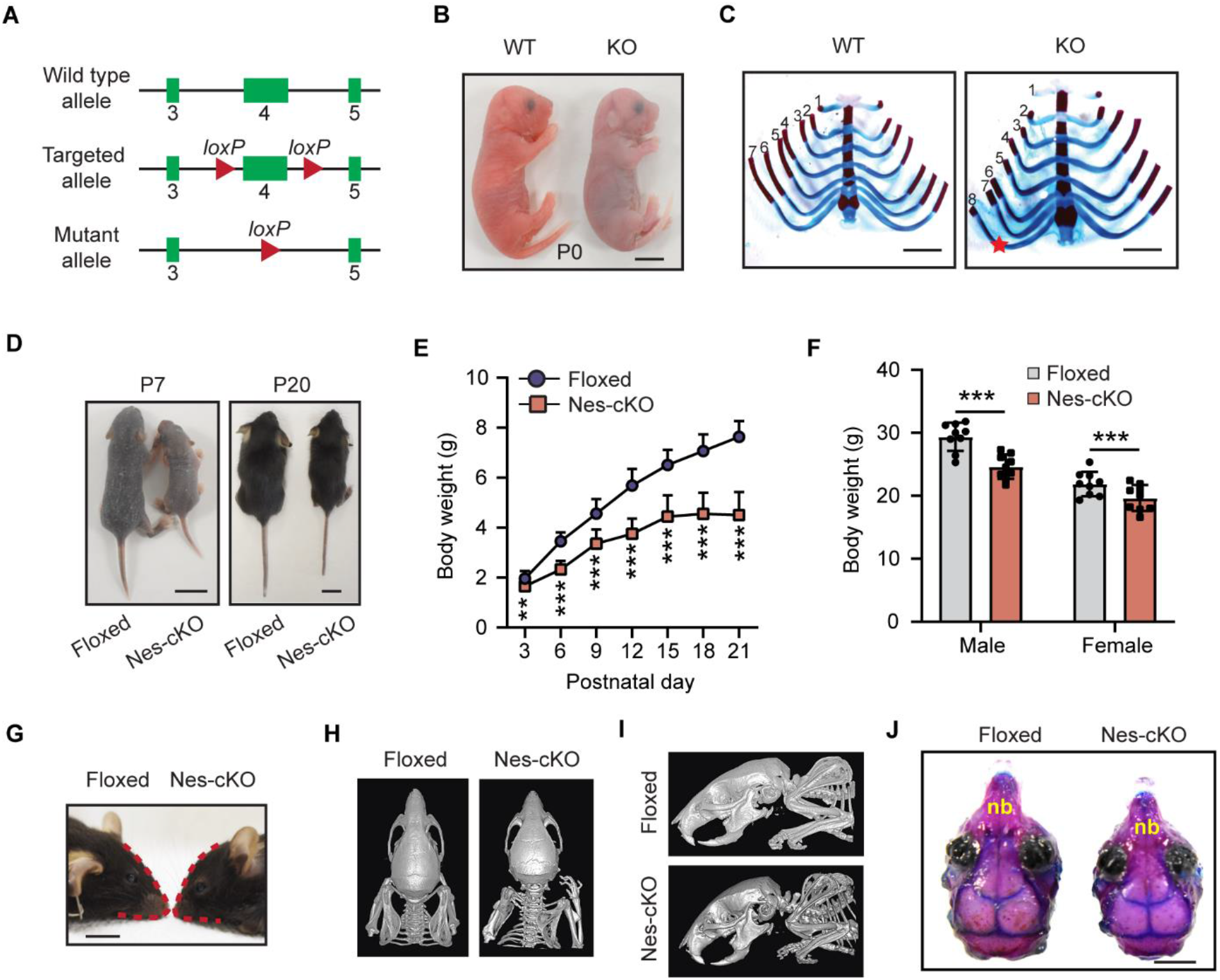
Characterization of *Ash1L* knockout mice. (**A**) Diagram showing the strategy for the generation of *Ash1L* conditional knockout mice. (**B**) Representative photos of wild-type and global *Ash1L*-KO newborns at P0. The *Ash1L*-KO newborns died at P0, bar = 5mm. (**C**) Photos showing the ventral view of rib cages of wild-type and *Ash1L*-KO mice. (**D**) Representative photos showing the body size of wild-type and *Ash1L*-Nes-cKO mice at P7 and P21, bar = 1cm. (**E**) Postnatal growth curve of wild-type and *Ash1L*-Nes-KO mice before weaning. Mixed gender body weight was plotted. For each group, n=15. *P* value calculated using a two-way ANOVA test. Error bars in graphs represent mean ± SEM. Note: \*\**P* < 0.01; \*\*\**P* < 0.001. (**F**) Body weight of adult wild-type and *Ash1L*-Nes-cKO mice. All the mice were measured at 3-month old. For each group, n=9. *P* value calculated using a two-way ANOVA test. Error bars in graphs represent mean ± SEM. Note: \*\*\**P* < 0.001. (**G**) Photo showing the craniofacial deformity of *Ash1L*-Nes-cKO mice, bar = 5mm (**H** and **I**) A dorsal (H) and lateral (I) view of mouse skull shown by micro-CT scanning. (**J**) A dorsal view of skull shown by bone staining. nb, nose bone, bar = 2mm.

To examine the function of *Ash1L* in the development of central nervous system, we deleted *Ash1L* in developing brains by crossing the *Ash1L-*cKO mice with a neural progenitor cell (NPC)-specific Cre (Nestin-Cre) mouse line. The Cre recombinase expressed in the NESTIN^+^ cells induced *Ash1L* deletion in NPCs as well as NPC-derived neuronal and glial lineages in embryonic developing brains. The mating (*Ash1L*^2f/2f^*;Nestin-Cre^−/−^* x *Ash1L*^2f/+^*;Nestin-Cre^+/-^*) produced wild-type (*Ash1L*^2f/2f^*;Nestin-Cre^−/−^*), heterozygous (*Ash1L*^2f/+^*;Nestin-Cre^+/-^*), and homozygous *Ash1L-* cKO (*Ash1L*-Nes-cKO, *Ash1L*^2f/2f^*;Nestin-Cre^+/-^*) progenies with the expected mendelian ratios. Compared to the wild-type littermates, the *Ash1L*-Nes-cKO newborns had similar body weight at birth and survived through early postnatal days. However, the *Ash1L*-Nes-cKO pups gradually displayed growth retardation, indicated by both smaller body size and lower body weight. The observed postnatal growth retardation appeared to be more drastic 2~3 weeks after birth, and the average body weight of *Ash1L*-Nes-cKO pups was approximately 50% less than that of wild-type littermates at P21 (Figure 1D-E). Although around 10% *Ash1L*-Nes-cKO pups died before weaning, the majority of surviving pups were able to grow to adulthood with their final body weight 5-10% lower than that of their wild-type adult littermates (Figure 1F). Similar to the craniofacial deformity observed in patients (*4, 9, 11, 12*), the adult *Ash1L*-Nes-cKO mice displayed an abnormal craniofacial appearance with a reduced eye-to-mouth distance, which was caused by shortened nose bones revealed by both micro-CT scan and skull bone staining (Figure 1G-J). No other obvious gross abnormalities of individual organs were observed in the *Ash1L*-Nes-cKO adult mice.

Next, we set out to investigate microscopic changes of *Ash1L-*KO mouse brains. Although the global *Ash1L-*KO newborns (P0) displayed normal gross brain appearance and size (Figure S2A-B), Nissl staining showed that the prefrontal cortex (PFC) of global *Ash1L-*KO mice was disorganized and lost its mini-columnar arrangement (Figure 2A), indicating malformations of cortical development (MCD). To further examine whether the disorganized PFC was caused by aberrant lamination during embryonic cortical development, we analyzed each cortical layer formation by immunostaining with cortical layer-specific antibodies. The results showed that the majority of layer II-III (L2/3)-specific SATB2^+^ neurons properly migrated to L2/3 in the wild-type PFC at P0. In contrast, some SATB2^+^ neurons in the *Ash1L*-KO PFC failed to migrate to the upper layers and scattered in the bottom layers (Figure 2B-C), suggesting that *Ash1L* deletion in developing brains resulted in delayed lamination of neuronal cells during embryonic cortical layer formation. To examine whether the *Ash1L* deletion in developing brains could lead to delayed myelination in postnatal brains as observed in some *ASH1L* mutation-related ASD/ID patients (*11*), we performed immunostaining for myelin basic protein (MBP) to examine the dynamic myelination in postnatal developing brains. The results showed that both wild-type and *Ash1L*-Nes-cKO cortices lacked discernable myelination at P0 (Figure 2D). However, compared to the wild-type controls that had increased myelination over time, the levels of myelination in the *Ash1L*-Nes-cKO cortex were significantly lower at P21 but reached to a comparable level around postnatal two months (Figure 2D-E), suggesting the *Ash1L* deletion in developing brains led to delayed myelination during early postnatal brain development.

**Figure 2.**
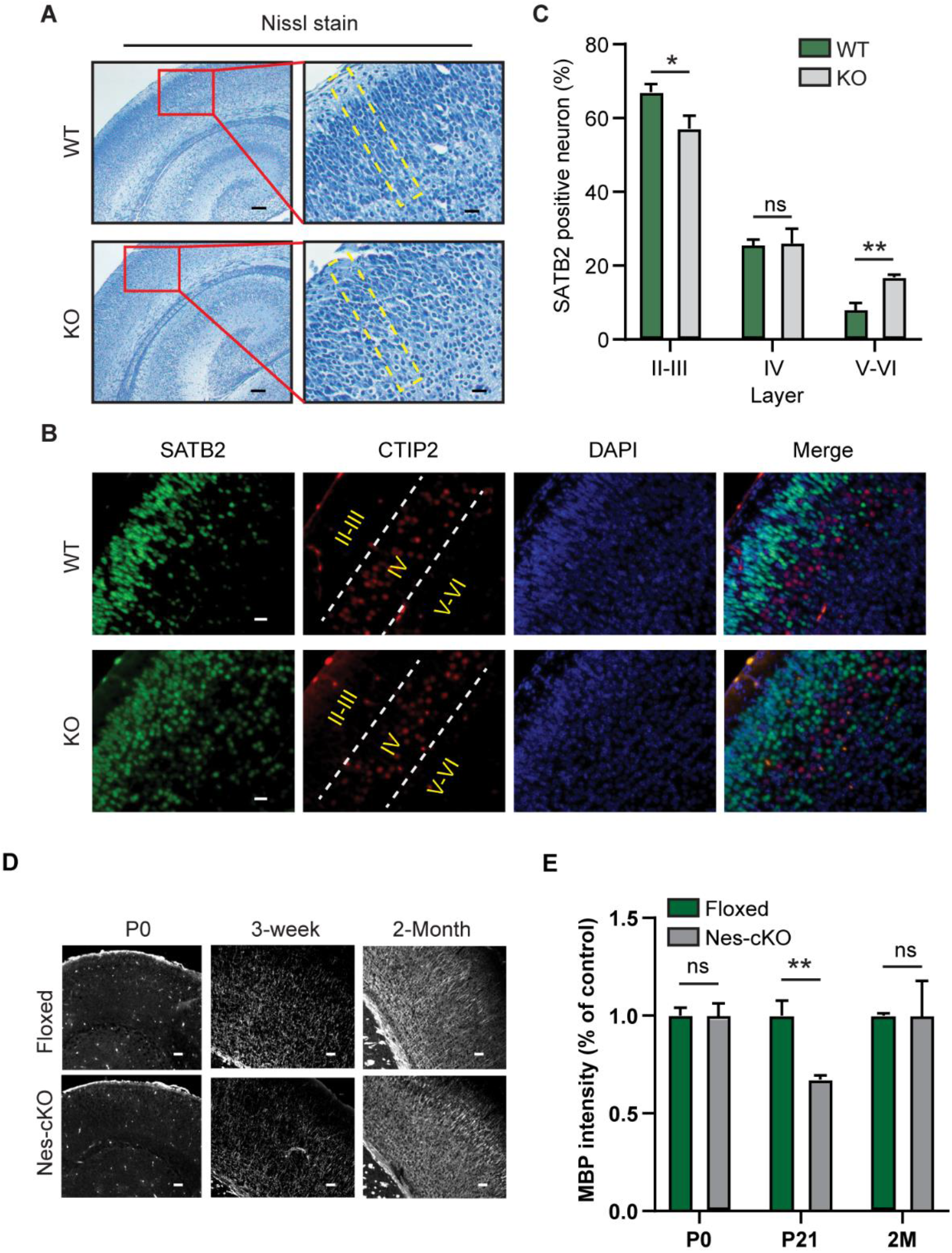
Abnormal brain development of *Ash1L* knockout mice. (**A**) Nissl staining showing the cortical histology of wild-type and global *Ash1L-*KO newborns. The mini-columnar arrangement of cortical cells is highlighted by yellow lines. Bar (left) = 100 μm, Bar (right) = 20 μm. (**B**) Photos showing the distribution of L2/3-specific SATB2^+^ cells and L4-specific CTIP2^+^ cells in wild-type and *Ash1L*-KO cortices at P0, bar = 20 μm. (**C**) Quantification of SATB2^+^ neurons in different layers. For each group, n=3. *P* value calculated using a two-tailed *t* test. Error bars in graphs represent mean ± SEM. Note: \**P* < 0.05; \*\**P* < 0.01; ns, not significant. (**D**) Photos showing the myelin basic protein (MBP) staining of P0, P21, and postnatal 2-month cortex, bar = 100 μm. (**E**) Quantitative MBP expression analyzed by the integrated fluorescence intensity in cortex. Values are percentage of control values ± SEM. For each group, n=3. *P* value calculated using a two-tailed *t* test. Note: **P < 0.01; ns, not significant.

After charactering the gross phenotypes of global and conditional *Ash1L* KO mice, we set out to investigate whether the *Ash1L* deletion in developing brains could lead to abnormal behaviors in adult mice. Since deficits in social interaction as well as repetitive and restricted behaviors are two major clinical manifestations found in human ASD patients, we first focused on testing these two core autistic behaviors. To this end, we performed a three-chamber test to examine the voluntary initiation of social interaction and the maintenance of interest in social novelty (*16*). The results showed that during the initiation of social interaction, the wild-type controls spent more time in the chamber containing a social partner, representing normal sociability. In contrast, the *Ash1L*-Nes-cKO mice displayed impaired initiation of social interaction, which was indicated by both reduced time and loss of preference for the chamber containing a social partner (Figure 3A-B). In the following social novelty tests, an unknown animal was introduced into the chamber and the subjects’ preference for social novelty was assessed. The wild-type mice displayed a preference for the novel animal, shown by the discrimination index favoring the chamber containing the novel mouse. In contrast, the *Ash1L*-Nes-cKO mice showed no preference for the chamber with the novel animal, indicating reduced social memory or lost interest in social novelty (Figure 3C-D). In addition to impaired social interaction, we observed that nearly all *Ash1L*-Nes-cKO adult mice displayed mild to severe front and hind paw clasping when suspended by tails, and 10-20% of mice had over-grooming that resulted in severe skin lesions (Figure 3E-F and Movie S1-2), suggesting the *Ash1L* deletion caused repetitive and compulsive behaviors, one of the core clinical manifestations observed in human ASD patients. Besides these two core autistic behaviors, some patients with *ASH1L* mutations had ID as their main clinical manifestation (*10–13*). Therefore, we further used the novel object recognition (NOR) test to examine whether the *Ash1L* gene deletion in developing brains could lead to impaired memory. 24 hours after the initial familiarization with two identical objects in an arena, the mice were allowed to explore the same arena in the presence of a familiar object and a replaced novel object (Figure 3G). The results showed that the wild-type mice spent more time to explore the novel object, reflecting normal recognition memory. In contrast, the *Ash1L*-Nes-cKO mice showed impaired recognition memory, indicated by the loss of preference to explore the novel object (Figure 3H). Furthermore, because some ASD/ID patients have elevated levels of anxiety, we examined anxiety-like behaviors by measuring time in the central areas of an open arena during habituation (Figure 3G). Compared to wild-type controls, although the *Ash1L*-Nes-cKO mice had increased locomotor activity, they spent about 50% less time exploring the arena center (Figure 3I-J), indicating the *Ash1L*-Nes-cKO mice had increased anxiety-like behaviors. Thus, the collective observations and behavior tests revealed that the loss of *Ash1L* in developing mouse brains resulted in both autistic-like behaviors and ID-like defects, which were featured by reduced sociability, loss of interest in social novelty, repetitive and compulsive behaviors, impaired recognition memory, and increased anxiety-like behaviors.

**Figure 3.**
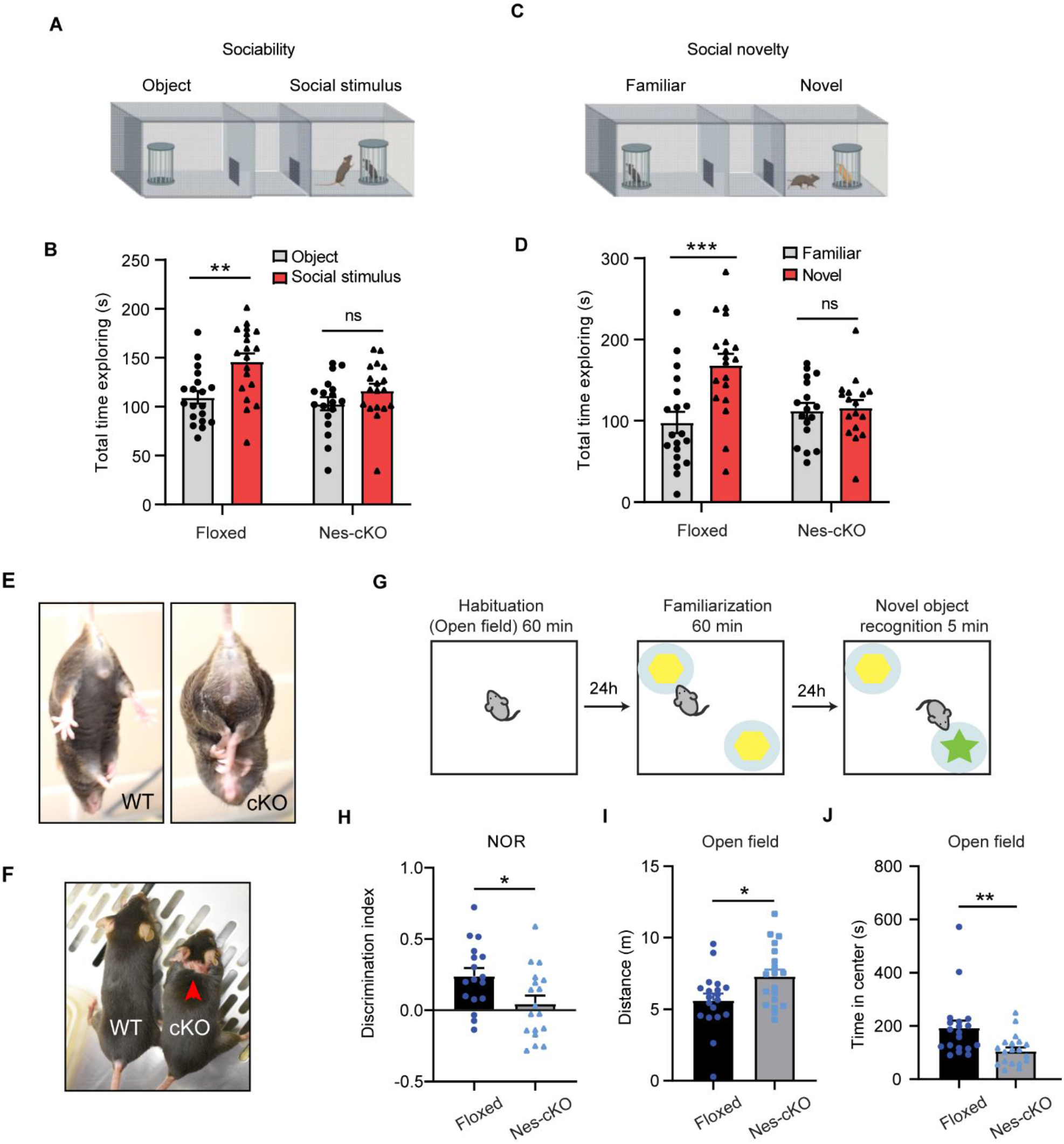
*Ash1L* knockout in developing brains causes autistic-like behavior and ID-like deficits. (**A**) 3-chamber tests for sociability. (**B**) Quantitative results showing the time wild-type (floxed) and *Ash1L*-Nes-cKO mice spent in the chamber containing a social partner in the 3-chamber sociability tests. (**C**) 3-chamer tests for social novelty. (**D**) Quantitative results showing the time wild-type (floxed) and *Ash1L*-Nes-cKO mice spent in the chamber containing a novel animal in the 3-chamber social novelty tests. For the 3-chamber sociability and social novelty tests (A-D), *Ash1L* wild-type mice, n=19; *Ash1L*-Nes-cKO mice, n=18. *P* value calculated using two-way ANOVA test. Error bars in graphs represent mean ± SEM. Note: \*\**P* < 0.01; \*\*\**P* < 0.001; ns, not significant. (**E**) Compared to wild-type mice (left panel), the *Ash1L*-Nes-cKO mice display paw clasping when suspended by tails (right panel). (**F**) Compared to wild-type mice, the *Ash1L*-Nes-cKO mice display skin lesions caused by over-grooming (red arrow). (**G**) Open field and novel object recognition (NOR) tests. (**H**) The quantitative discrimination ratio of NOR tests. The discrimination ratio was calculated as: (time spent on the novel object-time spent on the familiar object)/total time. *Ash1L*wild-type mice, n=17; *Ash1L*-Nes-cKO mice, n=18. *P* value calculated using a two-tailed *t* test. Error bars in graphs represent mean ± SEM. Note: \**P* < 0.05. (**I**) Total distance traveled in the open field arena. (**J**) Time spent in the center of the open field arena. For the open field tests (I-J), each group, n=19. *P* value calculated using a two-tailed *t* test. Error bars in graphs represent mean ± SEM. Note: \*\**P* < 0.01.

ASH1L is an epigenetic factor that regulates gene expression during development (*7*). To identify the genes regulated by ASH1L in developing brains, we performed RNA-sequencing (RNA-seq) analyses to examine differential gene expression between wild-type and *Ash1L*-deleted neural cells. To reduce the complexity in gene expression analyses on bulk brains containing multiple cell lineages and same cell lineages with different developmental stages, we set out to generate a tamoxifen-inducible *Ash1L-*cKO mouse line (*Ash1L*^2f/2f^*;Rosa26-CreER^T2+/+^*) by crossing the *Ash1L*-cKO mice with a Rosa26-CreER^T2^ line. The NPCs were isolated from the subventricular zone (SVZ) of brains and maintained in serum-free NPC culture medium. The deletion of *Ash1L* gene in the established NPCs was induced by 4-hydrotamoxifen (4OH-TAM) added in the medium for 10 days (Figure S3A). Quantitative reverse transcription PCR (qRT-PCR) and western blot (WB) analyses showed *Ash1L*/ASH1L expression reduced to less than 5% at mRNA and protein levels in the *Ash1L*-KO NPCs (Figure S3B-C). Both wild-type and *Ash1L-*KO NPCs were further induced to differentiate according to a well-established protocol (*17*), and immunostaining with lineage-specific markers was used to monitor the differentiation process. The results showed that both wild-type and *Ash1L-*KO cells expressed a NPC-specific marker NESTIN but not differentiation markers TUJ1 (neuron-specific tubulin III) or GFAP (glial fibrillary acidic protein) at day 0, indicating a homogenous NPC population before induced differentiation. 4 days after induced differentiation, both wild-type and *Ash1L*-KO cells had comparable decreased NESTIN^+^ NPCs and increased TUJ1^+^/GFAP^+^ differentiated neural cells (Figure S3D-G). To identify the differentially expressed genes in early NPC differentiation, we performed RNA-seq analyses 0, 12 and 24 hours after induced differentiation. The results identified total 2,475 upregulated and 2,808 downregulated genes during induced differentiation (cutoff: fold changes > 1.5, *p* < 0.01) (Figure S4H-I). Gene ontology (GO) enrichment analyses showed the upregulated genes had enriched GO terms involving nervous system development, while the downregulated genes involved metabolic processes and cell cycle regulation (cutoff: FDR < 0.05) (Figure S3J-K, Table S1, S2). Among all 2,475 genes upregulated in the wild-type NPC differentiation, 70 genes were found to have significantly reduced expression in the *Ash1L-*KO cells (cutoff: fold changes > 1.5, *p* < 0.01) (Figure 4A). These genes downregulated in the *Ash1L*-KO cells had enriched GO terms involving telencephalon development, regulation of cell communication, brain development, and central nervous development (cutoff: FDR < 0.05) (Figure 4B, Table S3), and Reactome pathway enrichment analysis revealed their involvement in postsynaptic signal transmission, including NMDA receptor activation (cutoff: FDR < 0.05) (Figure 4C, Table S4). Notably, multiple genes, such as *Emx2*, *Dbx2*, *Pcdh10*, and *Foxg1,* that were previously reported to play critical roles in normal brain development and related neurodevelopmental diseases (*18–25*) were found to have significantly reduced expression in the differentiating *Ash1L*-KO cells (cutoff: fold changes > 1.5, p < 0.01) (Figure 4D), suggesting that loss of ASH1L’s function in activating genes critical for normal brain development might be a possible molecular mechanism linking *ASH1L* mutations to neurodevelopmental defects of ASD/ID.

**Figure 4.**
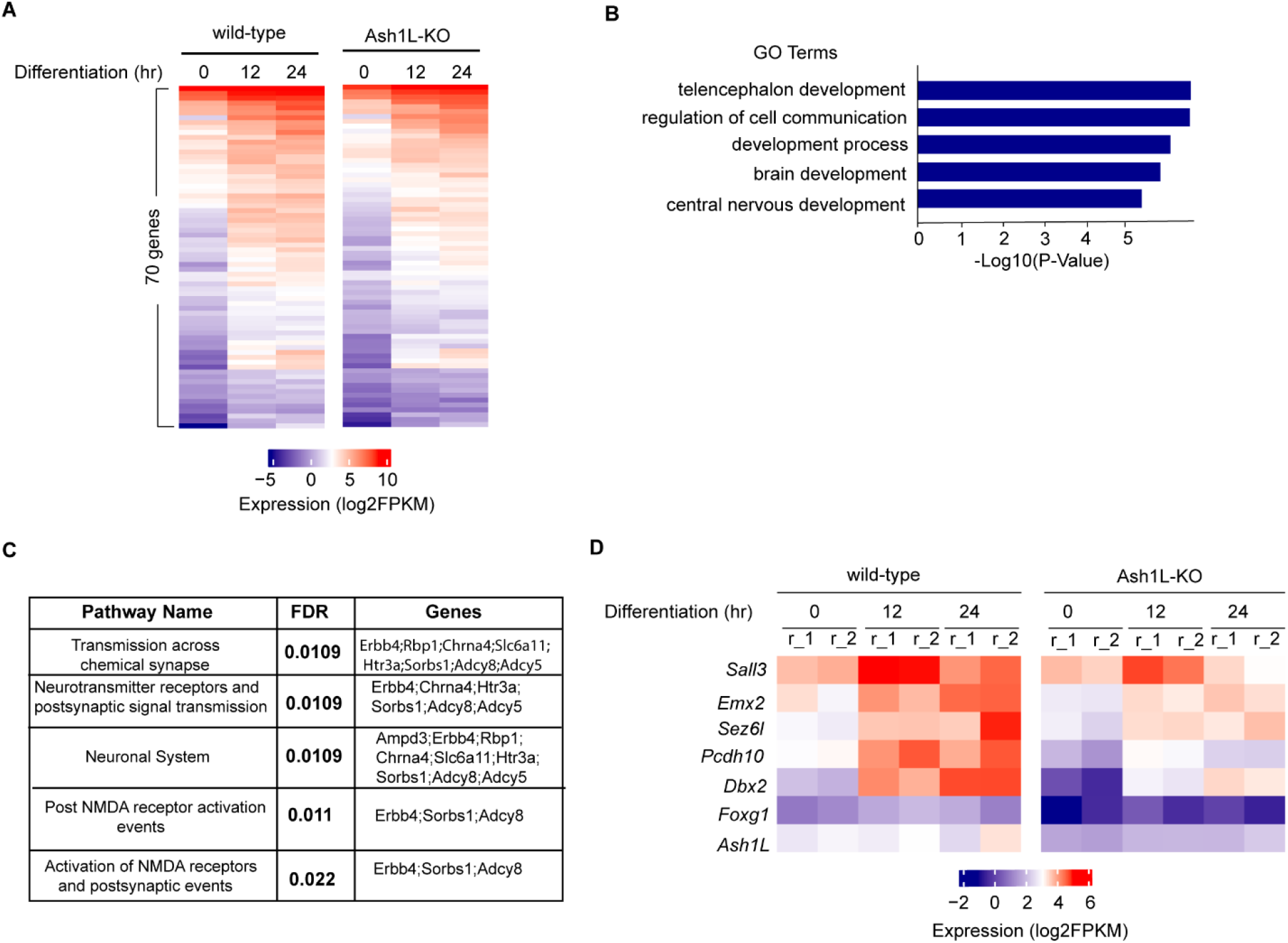
*Ash1L* knockout in neural progenitor cells impairs gene activation during induced differentiation. (**A**) Heatmap showing 70 genes upregulated in wild-type NPC differentiation have significantly reduced expression in the *Ash1L*-KO cells. (**B**) Gene ontology enrichment analysis showing the enriched GO terms of 70 downregulated genes in the *Ash1L*-KO cells during differentiation (FDR < 0.05). (**C**) Reactome pathway enrichment analysis showing the functional pathways related to the downregulated genes in the *Ash1L*-KO cells during differentiation. (cutoff: FDR < 0.05, gene membership ≥ 3). (**D**) Heatmap showing the representative downregulated genes in the *Ash1L*-KO cells during differentiation (cutoff: fold change >1.5, *p* < 0.01).

In this study, we used an animal model to demonstrate that the loss of *Ash1L* gene alone in developing mouse brains is sufficient to cause autistic-like behaviors and ID-like deficits in adult mice, which strongly suggests *ASH1L* gene mutations found in patients are likely to be the causative drivers leading to clinical ASD/ID (*9–14*). In addition, the early postnatal lethality found in the global *Ash1L-*KO newborns and the postnatal growth retardation observed in the *Ash1L*-Nes-cKO pups implicate *Ash1L* might also play important roles in establishing neural circuits in the developing hypothalamus, which is critical for normal feeding behaviors and early postnatal growth (*26*). Indeed, we observed the arcuate nucleus in hypothalamus is less developed in some global *Ash1L-*KO newborns (data not shown). It will be interesting to examine whether *ASH1L* mutations could cause similar hypothalamus dysfunctions that affect normal feeding behaviors and postnatal growth in human patients. Like the craniofacial deformity observed in the ASD/ID patients with *ASH1L* mutations, the *Ash1L*-Nes-cKO mice display craniofacial deformity with shortened nose bones. Since, in our animal model, *Ash1L* is also deleted in the NESTIN^+^ neural crest stem cells (NCSCs) that develop into craniofacial skeletal tissues (*27*), it is possible that the observed craniofacial deformity is caused by aberrant craniofacial skeletal formation from the *Ash1L*-deleted NCSCs. At the microscopic level, we observed both delayed lamination of cortical neurons and myelination formation during embryonic and early postnatal brain development, suggesting the loss of *Ash1L* in NPCs leads to delayed brain development in both neuronal and glial lineages. Biochemically, ASH1L is a histone H3 lysine 36-specific methyltransferase that facilitates gene expression through regulating transcriptional activation (*8*). Consistent with its biochemical functions, the RNA-seq analyses revealed the expression of multiple genes critical for normal brain development and highly related to human ASD are largely reduced in the differentiating *Ash1L-*KO NPCs. Of note, mutations of *FOXG1* gene, one of the genes downregulated in the *Ash1L-*KO cells, result in human FOXG1 syndrome. Interestingly, *FOXG1* syndrome and *ASH1L* mutations-induced ASD/ID have highly overlapping clinical manifestations (*24*), suggesting that ASH1L might function as a master regulator to activate the *FOXG1* and other critical genes during normal brain development, while mutations of *ASH1L* lead to mis-regulation of gene expression, disturbance of the normal brain developmental program, and brain functional abnormalities of ASD/ID. Finally, the new ASD/ID mouse model generated in this study recapitulates most clinical ASD/ID manifestations found in human patients, which provides an invaluable tool for further exploring the biological mechanisms underlying the pathogenesis of *ASH1L* mutations-induced ASD/ID, developing and testing new therapeutic approaches based on the functions of ASH1L and its regulated genetic pathways revealed by this study.

## Acknowledgements

We thank Drs. Jeremy Hix, Christiane Mallett, and Erik Shapiro for micro-CT scan imaging assays. MSU genomics core facility processed the next-generation sequencing. This work was supported by the National Institutes of Health (grant R01GM127431).

## Author contributions

J.H. conceived the project. Y.G., N.D., J.H. performed the experiments. Y.G. and M.B.A. maintained the mouse colonies. Y.W. prepared the recombinant protein for the Ash1L antibody generation. A.J.R. oversaw the mouse behavior tests. G.I.M. and J.H. performed the sequencing data analysis. Y.G., N.D., A.J.R., and J.H. interpreted the data. Y.G., N.D., and J.H. wrote the manuscript.

## Supplementary Materials

Table S1. Result of gene ontology enrichment analysis on the upregulated genes in the wild-type NPCs during induced differentiation.xlsx

Table S2. Result of gene ontology enrichment analysis on the downregulated genes in the wild-type NPC during induced differentiation.xlsx

Table S3. Result of gene ontology enrichment analysis on the reduced expressed genes in the *Ash1L*-KO NPCs during induced differentiation.xlsx

Table S4. Result of Reactome pathway enrichment analysis on the reduced expressed genes in the *Ash1L*-KO NPCs during induced differentiation.xlsx

Table S5. Sequences of all primers used in this study.xlsx

Video S1. Posture of wild-type mice upon tail suspension

Video S2. Paw clasping of *Ash1L*-Nes-cKO mice upon tail suspension

## Materials and Methods

### Mice

The *Ash1L* conditional knockout target construct was generated by modifying the BAC clone (RP24-394C15) based on the Recombneering method (*28*). Two LoxP elements were inserted into the exon 4-flanking sites. The targeting construct was electroporated into the C57BL/6:129 hybrid murine ES cells. Homologous recombinant ES cell clones were identified PCR-based genotyping and injected into blastocysts. The genetic modified ES cells were micro-injected to blastocysts and transferred to the uterus of CD-1 pseudo-pregnant females to generate chimeric founder mice. The chimeric founder mice were crossed to a FLP recombinase mouse line to remove the FRT flanked selection cassette. The *Ash1l* global knockout mice were generated by mating *Ash1L* floxed mice with CMV-cre mice (B6.C-Tg (CMV-cre) 1Cgn/J, The Jackson Laboratory). The *Ash1L* neural conditional knockout mice were generated by mating *Ash1L* floxed mice with Nestin-cre mice (B6.Cg-Tg (Nes-cre) 1Kln/J, The Jackson Laboratory). The 4-hydrotamoxifen inducible *Ash1L*-cKO mice were generated by mating *Ash1L* floxed mice with Rosa26-CreER^T2^ mice (B6.129-*Gt(ROSA)26Sor^tm1(cre/ERT2)Tyj^*/J, The Jackson Laboratory). All mice were backcrossed to C57BL/6 mice for at least five generations to reach a pure C57BL/6 background before experiments. Mice were housed under standard conditions (12h light: 12h dark cycles) with food and water ad libitum. The data obtained from all embryos were pooled without discrimination of sexes for the analysis. All mouse experiments were performed with the approval of the Michigan State University Institutional Animal Care & Use Committee.

### Genotyping

Genomic DNA was extracted from mouse tails with lysis buffer of 0.01M NaOH. After neutralization with Tris-HCl (PH 7.6), the extracted genomic DNA was used for genotyping PCR assays. Primers used for genotyping were listed in table S5.

### Isolation, culture and induced differentiation of neural progenitor cells

The neural tissues were isolated from the subventricular zone of P30 brains and dissociated with 0.05% trypsin-EDTA (Life Technologies) at 37 °C for 15 minutes. The reaction was stopped by trypsin inhibitor (10mg/ml, Worthington Biochemical Corporation). The dissociated cells were washed with cold PBS for 3 times and plated onto the non-coated petri dishes in the Neurobasal medium (Life Technologies) that was supplemented with 1x B27 supplement (Gibco), 1x GlutaMAX (Life Technologies), 20 ng/ml murine epithelial growth factor (Peprotech), 20 ng/ml basic fibroblast growth factor (Peprotech), and 100U/ml penicillin/streptomycin (Life Technologies). 5 ~ 7 days later, the neurospheres formed by proliferating NPCs were collected and re-plated onto the Poly-L-Ornithine (R&D systems) and Laminin (Corning)-coated plates to form monolayer culture. The neural progenitor cells were passaged at 1:5 ratio every 3 days. To induce NPCs differentiation, the NPC monolayer cells were gently washed with PBS for 3 times and cultured under the Neurobasal medium supplemented with 1x N2 supplement (Giboco), 1x GlutaMAX (Life Technologies).

### Induced *Ash1L* deletion in neural progenitor cells

*Ash1L* gene deletion in the *Ash1L*^2f/2f^; Rosa26-CreER^T2+/-^ NPCs were induced by addition of 4-hydrotamoxifen (Sigma-Aldrich) at 0.1 μM in the culture medium. Genotyping were used to confirm the Ash1L gene deletion 10 days after 4-hydrotamoxifen treatment. The exclude any effects caused by 4-hydrotamoxifen, the confirmed *Ash1L*-deleted NPCs were further cultured without 4-hydrotamoxifen for at least 3 passages before further experiments.

### Alizarin red/alcian blue bone staining

Whole mouse carcasses were collected after euthanasia, defatted for 2–3 days in acetone, stained sequentially with Alcian blue and alizarin red S in 2% KOH, cleared with 1% KOH/20% glycerol, and stored in 50% EtOH/50% glycerol.

### Microcomputed tomography (micro-CT)

Skulls were serially imaged using a PerkinElmer Quantum GX micro-CT scanner. The following image acquisition parameters were used at each scan time point: 4 minutes acquisition; 90 kVp/88 μA; Field of View (FOV), 45 mm; pixel resolution, 90 μm. Then, tissue sections were collected for histology and Ta analysis using ICP-OES.

### Nissl staining

The paraffin blocks were prepared by Division of Human Pathology of Michigan State University. Briefly, mouse brain sections were dewaxed in xylene and rehydrated in alcohol. Then, sections were stained in toluidine buffer [1g toluidine blue (Sigma) in 100 mL 95% ethanol] at room temperature for 20 min. Quick rinse in tap water and 70% ethanol to remove excess stain. After dehydration and wax, sections were mounted in mounting media H5000 (Vector Laboratories). Images were captured using a Zeiss Axio Imager microscope (Carl Zeiss GmbH, Oberkochen, Germany) and an installed AxioCam HRc camera (Carl Zeiss GmbH) with image acquisition via Zeiss Zen Pro software (v.2.3; Carl Zeiss GmbH).

### Immunostaining

Mouse tissues were fixed in 4% PFA in PBS overnight at 4 °C and embedded in paraffin. For immunofluorescence, tissue sections of 5 μm were cut, dewaxed and rehydrated. Antigen retrieval was performed by microwaving the sections on 0.01M sodium citrate buffer (pH 6.0) for 4 min. Tissue sections were blocked in 5% normal donkey serum (NDS) for 30 min after sensing with PBS. Tissue sections then were incubated with primary antibodies diluted in 5% NDS overnight at 4 °C. Antibodies used were: mouse anti-SATB2 (1:10, ab51502, abcam), rat anti-CTIP2 (1:100, ab18465, abcam), and rabbit anti-MBP (1:500; 78896; Cell Signaling technology). After washing with PBS, sections were incubated with Alexa Fluor 488 donkey anti-mouse IgG (1:300; 715-545-150; Jackson ImmunoResearch) or R-Phycoerythrin AffiniPure F(ab’)₂ Fragment Donkey Anti-Rat IgG (1:300; 712-116-153; Jackson ImmunoResearch) for 1 h and mounted using Vectorshield mounting media with DAPI (H1200, Vector Laboratories). Images were captured using a Zeiss Axio Imager microscope (Carl Zeiss GmbH, Oberkochen, Germany) and an installed AxioCam HRc camera (Carl Zeiss GmbH) with image acquisition via Zeiss Zen Pro software (v.2.3; Carl Zeiss GmbH).

### ASH1L antibody generation

The Polyclonal Rabbit anti-ASH1L antibody was generated by Pocono Rabbit Farm & Laboratory. The recombinant mouse ASH1L peptides (aa 2053-2347) were used as antigen. The antibodies were purified by immunoaffinity chromatography using antigen-coupled affi-gel 10 (Bio-rad).

### Western Blot analysis

Total proteins were extracted by RIPA buffer and separated by electrophoresis by 8-10% PAGE gel. The protein was transferred to the nitrocellulose membrane and blotted with primary antibodies. The antibodies used for Western Blot and IP-Western Blot analyses included: rabbit anti-Ash1L (1:1000, in house) and IRDye 680 donkey anti-rabbit second antibody (1: 10000, Li-Cor). The images were developed by Odyssey Li-Cor Imager (Li-Cor).

### qRT-PCR assays

RNA was extracted and purified from cells using QI shredder (Qiagen) and RNeasy (Qiagen) spin columns. Total RNA (1 μg) was subjected to reverse transcription using Iscript reverse transcription supermix (Bio-Rad). cDNA levels were assayed by real-time PCR using iTaq universal SYBR green supermix (Bio-Rad) and detected by CFX386 Touch Real-Time PCR detection system (Bio-Rad). Primer sequences for qPCR are listed in Supplementary Table 4.

### RNA-seq sample preparation for HiSeq4000 sequencing

RNA was extracted and purified from cells using QI shredder (Qiagen) and RNeasy (Qiagen) spin columns. Total RNA (1 μg) was used to generate RNA-seq library using NEBNext Ultra Directional RNA library Prep Kit for Illumina (New England BioLabs, Inc) according to the manufacturer’s instructions. Adapter-ligated cDNA was amplified by PCR and followed by size selection using agarose gel electrophoresis. The DNA was purified using Qiaquick gel extraction kit (Qiagen) and quantified both with an Agilent Bioanalyzer and Invitrogen Qubit. The libraries were diluted to a working concentration of 10nM prior to sequencing. Sequencing on an Illumina HiSeq4000 instrument was carried out by the Genomics Core Facility at Michigan State University.

### RNA-Seq data analysis

RNA-Seq data analysis was performed essentially as described previously. All sequencing reads were mapped mm9 of the mouse genome using Tophat2 (*29*). The mapped reads were normalized to reads as Reads Per Kilobase of transcript per Million mapped reads (RPKM). The differential gene expression was calculated by Cuffdiff program and the statistic cutoff for identification of differential gene expression is p < 0.01 and 1.5-fold RPKM change between samples. The heatmap and plot of gene expression were generated using plotHeatmap and plotProfile in the deepTools program (*30*). The differential expressed gene lists were input into the GENEONTOLOGY for the GO enrichment analyses (http://geneontology.org/). Pathway enrichment analysis was carried out using PyIOmica’s interface for Reactome (*31, 32*), with false discovery rate, FDR < 0.05, and a filter with a minimum of 3 genes being identified a pathway.

### Open field test

The open-field apparatus consisted of a custom-made, square white polyvinylchloride foam box (38 x 38 x 35cm). Their behavior was recorded for the first 10 min of habituation to measure time spent in open field, time spent in corners, and time freezing with a digital CCD camera connected to a computer running an automated video tracking software package (Clever Sys).

### Novel object recognition test (NOR)

NOR was assessed using a 3-day paradigm that included habituation, training, and testing as described previously (*33*). Each day, mice were acclimated for 60 min to the behavioral testing room before assessment. All testing was performed under red lights. During habituation (day1), mice were placed into the open field apparatus, a square white polyvinylchloride foam 38×38×35 cm box, for 60 min. For training (day 2), mice were placed in the same apparatus, where two similar objects had been placed near the corners. The object pairs consisted of miniature wheels, knobs, spark plugs, and Lego blocks. Mice were allowed to freely explore the objects for 30 min. They did not show consistent biases for any one object over another. For testing (day 3), mice were placed in the same apparatus, but this time one object of the pair was replaced with another dissimilar object (novel object), and they were allowed to freely explore for 5 minutes. Their behavior was recorded, and the time the mice spent with their nose oriented towards the object within a 3.5cm radius of the object was considered as exploration time. Throughout testing, objects and apparatus were cleaned with 70% ethanol between trials. Discrimination index was calculated as:

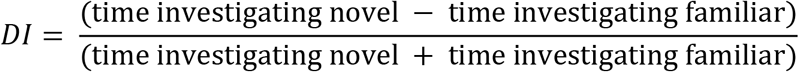

### Sociability and preference for social novelty test

This test was adapted from Crawley’s sociability and preference for social novelty protocol (*16*), which consists of three phases. Before testing, mice were acclimated for 60 min to the behavioral testing room under red lights. In phase 1 (habituation), the experimental mouse was placed in the center of a three-chamber apparatus (polyvinylchloride, 60×40×22 cm, fig.1) and allowed to freely explore for 5 minutes. During this time, the mouse had free access to all three chambers, which are connected by small openings at the bottom of the dividers. In phase 2 (sociability), two identical, wire cup-like containers were placed one in each side chamber. In this phase, an unfamiliar same-sex mouse was placed in one of the containers (“social stimulus”), while the other remained empty (“object”). The experimental mouse was allowed to freely explore the three chambers again for 5 minutes. In phase 3 (social memory), the container with the mouse (now “known”) was moved to the opposite chamber, and a new same-sex mouse (“unknown”) was placed in the other container. The experimental mouse was allowed to freely explore the three chambers for 5 minutes. During the three phases, the behavior spent in each of the chambers was recorded. Throughout testing, objects and apparatus were cleaned with 70% ethanol between trials.

### Statistical analysis

All statistical analyses were performed using GraphPad Prism 8 (GraphPad Software). Parametric data were analyzed by a two-tailed *t* test or two-way ANOVA test for comparisons of multiple samples. *P* values < 0.05 were considered statistically significant. Data are presented as mean ± SEM.

#### Supplemental figures

**Figure S1.**
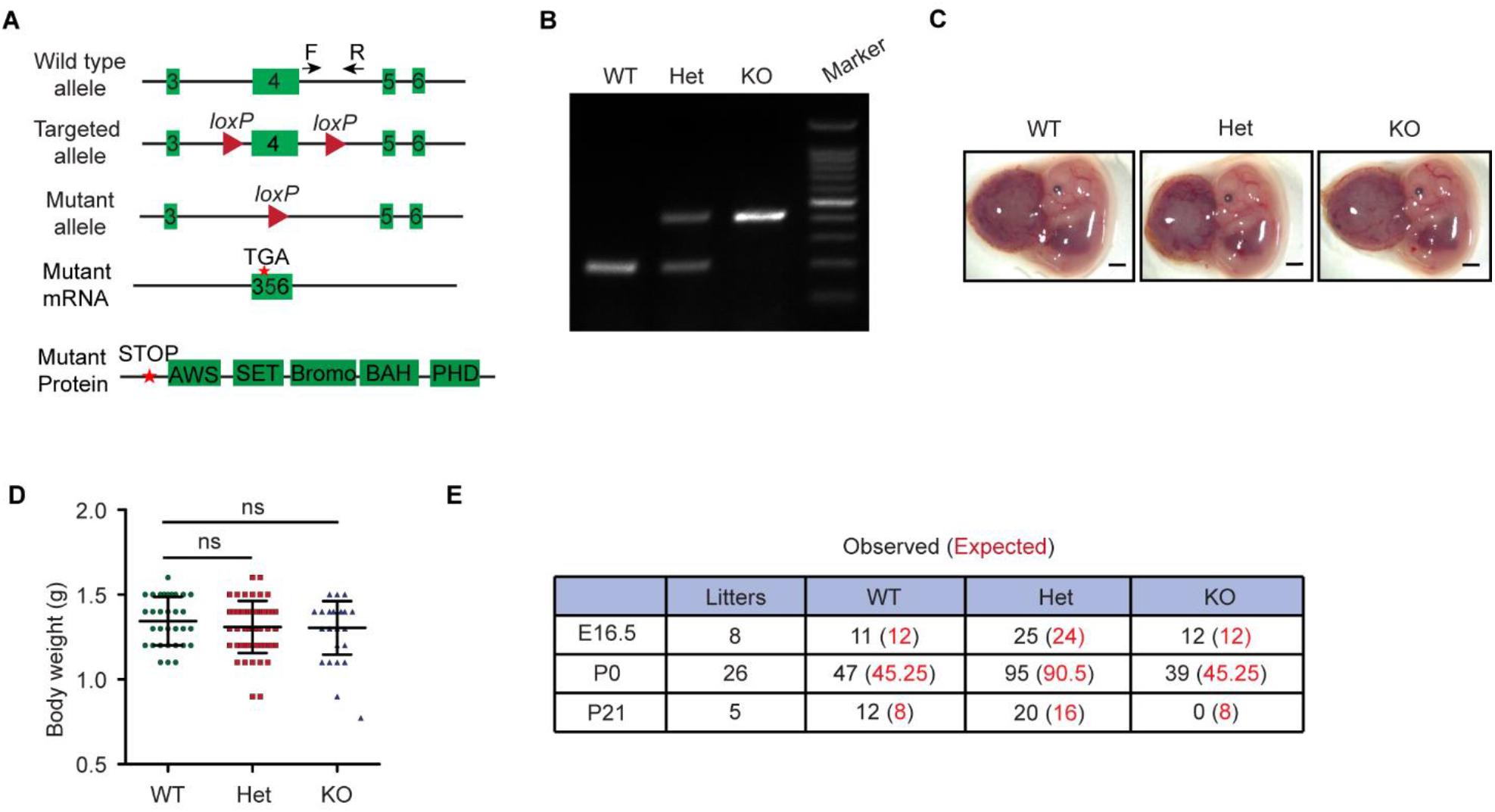
Characterization of *Ash1L* knockout mice. (**A**) Diagram showing the strategy for the generation of *Ash1L* conditional knockout mice. Cre-mediated deletion of exon 4 results in an altered spliced mRNA with a premature stop codon, which generates a truncated protein without all functional AWS, SET, Bromo, BAH and PHD domains. The arrows labeled as F and R represent the genotyping primers. (**B**) Genotyping results showing the PCR results of wild-type, heterozygous, and homozygous *Ash1L* knockout mice. (**C**) Photos showing the gross morphology of wild-type, heterozygous, and homozygous Ash1L knockout embryos and placentas at E13.5, bar = 500 μm. (**D**) Body weight of global *Ash1L-*KO mice at P0. For Ash1L WT, n=32; for *Ash1L* Het, n=53; for *Ash1L* KO, n=23. *P* value calculated using a two-tailed *t* test. Error bars in graphs represent mean ± SEM. Note: ns, not significant. (**E**) The genotyping results of global *Ash1L*-KO mice analyzed at embryonic and postnatal stages.

**Figure S2.**
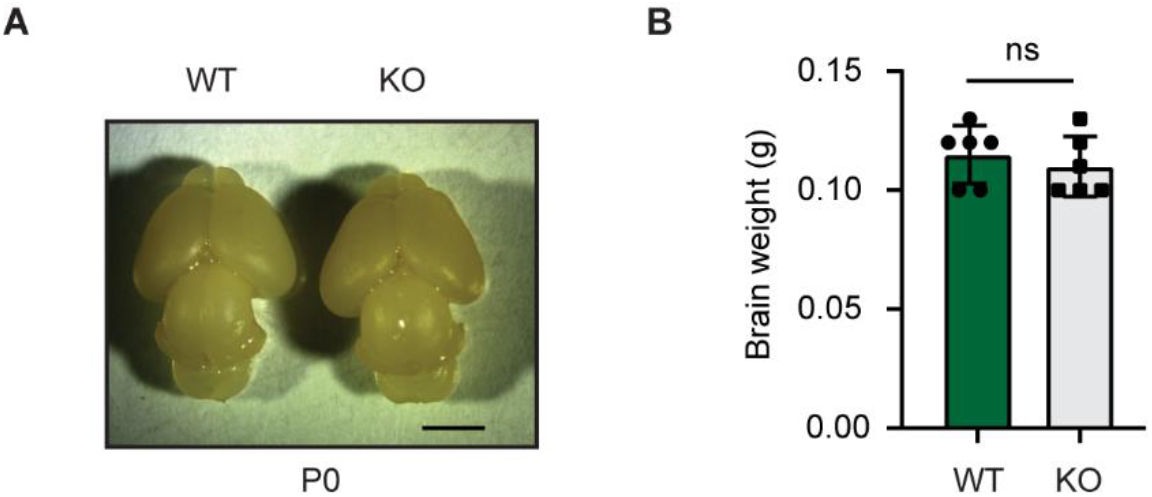
Abnormal brain development of *Ash1L* knockout mice. (**A**) Photo showing the gross brain morphology of wild-type and global *Ash1L-*KO mice at P0, bar = 2 mm. (**B**) The brain weight of P0 wild-type and global *Ash1L*-KO mice. For each group, n=6. *P* value calculated using a two-tailed *t* test. Error bars in graphs represent mean ± SEM. Note: ns, not significant.

**Figure S3.**
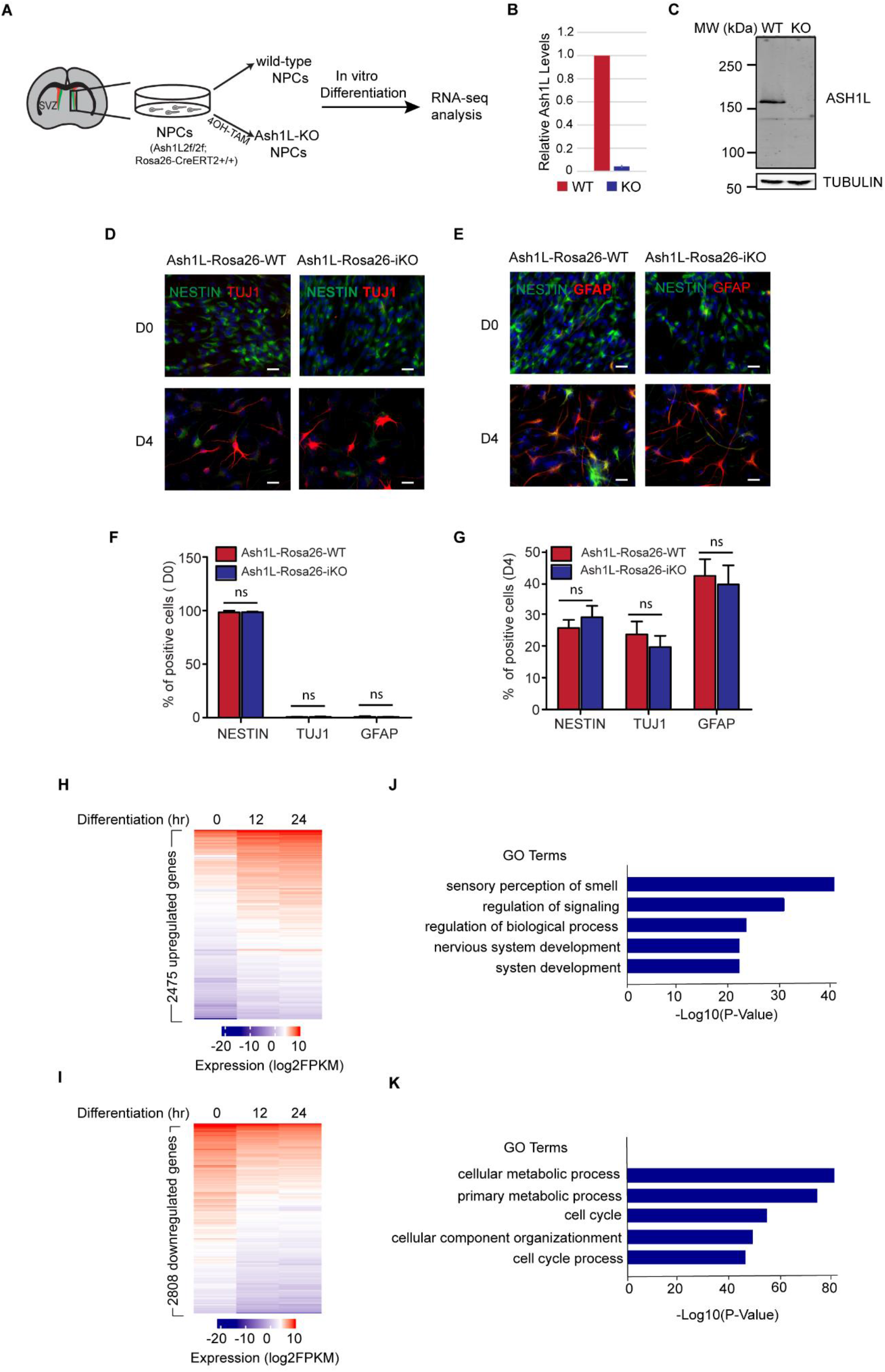
*Ash1L* knockout in neural progenitor cells impairs gene activation during induced differentiation. (**A**) Flowchart showing the experimental design for gene expression analysis. (**B**) qRT-PCR analysis showing the expression levels of *Ash1L* in wild-type and knockout NPCs. The results were normalized against levels of *Gapdh* and the expression level in wild-type NPC was arbitrarily set to 1. The error bars represent the standard deviation (n=3). (**C**) Western blot analysis showing the protein levels of ASH1L in wild-type and knockout NPCs. (**D-E**) Photos showing the NESTIN^+^/TUJ1^+^ (D) and NESTIN^+^/GFAP^+^ cells (E) at D0 and D4 after induced differentiation, bar = 20 μm. (**F-G**) Quantification of NESTIN^+^, TUJ1^+^, and GFAP^+^ cells at D0 (F) and D4 (G) after induced differentiation. For each group, n=3. *P* value calculated using a two-tailed *t* test. Error bars in graphs represent mean ± SEM. Note: \**P* < 0.05; \*\**P* < 0.01; ns, not significant. (**H**) Heatmap showing 2,475 upregulated genes in wild-type NPC differentiation (cutoff: change folds > 1.5, p < 0.01). (**I**) Heatmap showing 2,808 downregulated genes in wild-type NPC differentiation (cutoff: change folds > 1.5, p < 0.01). (**J**) Gene ontology enrichment analysis showing the enriched GO terms of 2,475 upregulated genes in the wild-type NPC differentiation (FDR < 0.05). (**K**) Gene ontology enrichment analysis showing the enriched GO terms of 2,808 downregulated genes in the wild-type NPC differentiation (FDR < 0.05).

